# Kinetic Model of Translational Autoregulation

**DOI:** 10.1101/432658

**Authors:** Vivian Tyng, Michael E. Kellman

## Abstract

We investigate dynamics of a kinetic model of inhibitory autoregulation as exemplified when a protein inhibits its own production by interfering with its messenger RNA, known in molecular biology as translational autoregulation. We first show how linear models without feedback set the stage with a nonequilibrium steady state that constitutes the target of the regulation. However, regulation in the simple linear model is far from optimal. The negative feedback mechanism whereby the protein “jams” the mRNA greatly enhances the effectiveness of the control, with response to perturbation that is targeted, rapid, and metabolically efficient. Understanding the full dynamics of the system phase space is essential to understanding the autoregulation process.

## Introduction

Autoregulation is extremely important in a multitude of contexts. Examples range from the molecular level of gene regulation, to organ level control of physiological processes, e.g. control of blood flow under blood pressure variation.^1^ Perhaps the most common type of autoregulation is negative (or repressive or inhibitory) regulation in an activator-repressor network,^2^ represented as follows in a very simple network diagram shown in Fig. 1: with the arrow conventionally representing activation and the block repression or negative feedback.

**Figure 1.**
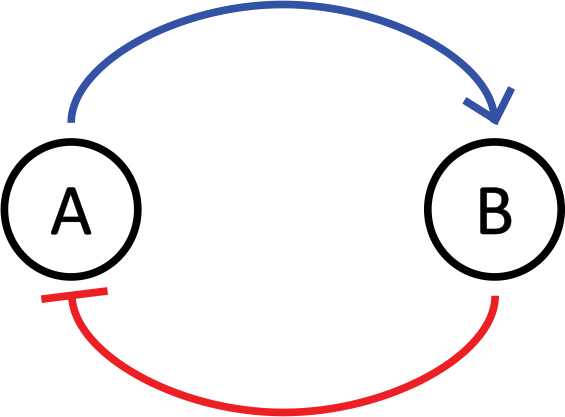
Schematic activator-repressor network.

A very important example in molecular biology is when a protein regulates its own production by inhibiting the translation of its gene into mRNA. This is *transcriptional autoregulation*. In another important mechanism, the subject of the investigation here, the control takes place through binding of the protein to the mRNA produced by the gene for the production of the protein. This mechanism wherein the protein “jams” its own mRNA template is called *translational autoregulation*. These epigenetic mechanisms of autoregulation take place among the elements of the “central dogma” of molecular biology that “DNA makes RNA makes Protein.” A schematic is shown in Fig.2. Examples of translational autoregulation, which has proven to be a widespread phenomenon,^3^range from the gp32 protein^4–7^ involved in the DNA replication process of T4 virus in *E. coli*, to thymidylate synthase (TS), a protein that plays important roles in a variety of common cancers.^8^

**Figure 2:**
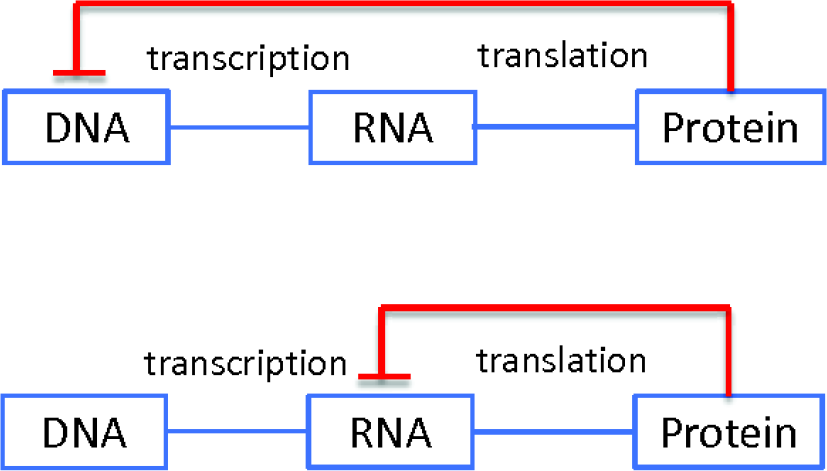
Transcriptional and translational autoregulation of gene expression.

## Kinetic model for translational autoregulation with negative feedback

In this section we consider the motivation for detailed kinetic analysis of autoregulation, then describe the elements of the specific model of translational autoregulation that is the focus of this paper.

### Why a kinetic model?

Translational autoregulation is an example of a process in a far-from equilibrium system, and as such has intrinsic interest as a problem in kinetics. Moreover, kinetic analysis is recognized as an essential element for understanding of biologically crucial gene regulation processes.^9–11^ Protein and RNA kinetics are becoming more accessible with advances in proteomics and transcriptomics.^12^ We seek to advance understanding of the protein-mRNA regulatory and kinetic problem by adding to earlier models of this system^11^ the additional fundamental element of nonlinear feedback – a combination which has not to our knowledge been exploited in the mathematical analysis of translational autoregulation kinetics, though feedback is most certainly known as an essential element of mathematical systems biology.^9,10^

A kinetic model gives the *sequence in time* of the concentrations of all the species in the regulatory system.^13^ Here we will be studying the full dynamics of a two-component model. A simple qualitative characterization like the network diagram in Fig. 1 simply cannot do full justice to the regulatory process. The purpose of our investigation of a kinetic model of autoregulation is to understand how the detailed quantitative dynamics of the system of rate constants with negative feedback and cooperative behavior sets the stage for this particular type of autoregulation, and optimizes its behavior.

We will confine our attention here to the translational autoregulatory process. The DNA transcriptional mechanism, already considered by Rosenfeld et al.^14^ but in a lower-dimensional approach than we adopt, has other features that deserve a separate treatment. Others^11,12^ have considered two-component models of translational regulation that do not however include feedback. These early models are formally identical to the “linear” models that will be our starting point here. Hargrove and Schmidt^11^ argue as we do for the great importance of these simple models for beginning to understand gene regulatory processes at the mathematical systems level. Among our central goals are to see how the phase space view of dynamics adds greatly to understanding of even the simple linear systems. Ref.^11^ considered the individual concentration changes vs. time for specific initial conditions, but did not plot the two-dimensional dynamics in a phase space “portrait” in the vicinity of the steady state, i.e. the target of the negative autoregulation. We find that this helps greatly to understand the linear systems, and then to see how the addition of feedback contributes immensely to the possibilities for optimizing the autoregulatory control.

### Elements of an effective control scheme

To build an effective control scheme, we need a system in which the control is *targeted* and *rapid*. We may also want to take into account the *metabolic energy demands* of various possible kinetic schemes (including their parameters) on an organism, aiming for control that is *efficient*. To achieve all of this, we need two elements, as shown in Fig. 3: (1) a basic “barebones” linear kinetic scheme of reactions without feedback. This has rates for production (via gene transcription) and degradation of the mRNA; and production (via translation of the mRNA by the ribosome machinery) and degradation of the protein. As emphasized quite some time ago by Hargrove and Schmidt,^11^ this is already a simple regulatory system that has the feature of a unique steady state that is the basic target of the regulation, which dampens deviations from the steady state. (2) Then, we need additional features of feedback and cooperativity (terms defined mathematically below) that give to the autoregulation the robust aspects of control outlined above (“targeted, rapid, efficient“). The feedback and cooperativity “tune” the barebones linear network to give superior performance. Fig. 3 shows the autoregulatory network with the species mRNA (*m*) and protein (*P*), with arrows representing the barebones linear scheme, and the negative feedback with cooperativity depicted as the blocking by *P* of its own production with *m*.

**Figure 3:**
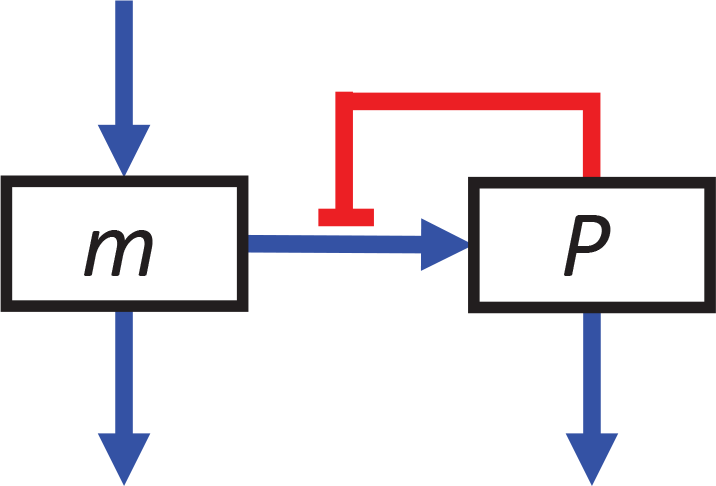
Schematic of the translational autoregulatory model including feedback. m: mRNA, P: protein.

In the following sections, we will build the barebones linear model and examine its essential features and limitations of performance, then tune this model with feedback and cooperativity to see how this gives superior regulation. We will see that each component of the system affects the performance and the tuning of the other components. Our plan is the following. First, in Section we imagine a cell or virus that seeks a certain steady state level of the protein *P*. We construct a linear model that gives the desired concentration. We will see that there is a natural classification of the dynamics into three qualitatively different types in the linear model, depending on a key ratio of two of the parameters. Then, in Section we imagine the cell or organism adapting by adding feedback and cooperativity to get better autoregulatory control (e.g. faster response times) than is afforded by the linear model, while maintaining the same target steady state level of *P*. We will choose a numerical “feedback strength,” and adjust the other parameters accordingly. We will compare the behavior of each of the three types of control dynamics with feedback with that of the corresponding linear system. Then, in Section we make a systematic comparison of the autoregulatory control performance of the linear systems and the corresponding systems with feedback.

## Linear model with steady state

We imagine a cell or virus that seeks a certain steady state level *P_SS_* of the protein, and systematically construct a linear model, without feedback, that would accomplish this goal.

This “barebones” linear model already represents the simplest case of a control scheme. Great insight can be obtained because of its simplicity. It needs to be stated that the linear model has been considered long ago and its essential importance in protein regulation recognized; our equations (1-2) are identical in content to the system of Ref.^11^ However, our phase space portraits with organization into three classes are new, as is our addition later of nonlinear feedback as an essential element. Another difference is that Ref.^11^ emphasizes analytical solutions for the linear model, while we also emphasize numerically computed dynamics for both the linear system and the system with nonlinear feedback, as is essential for the latter. As noted already, we find that there are three qualitatively different types of linear models, depending on the chosen parameters. We will examine the control properties of each of these versions of the model, with a view toward their individual advantages and shortcomings.

**Figure 4:**
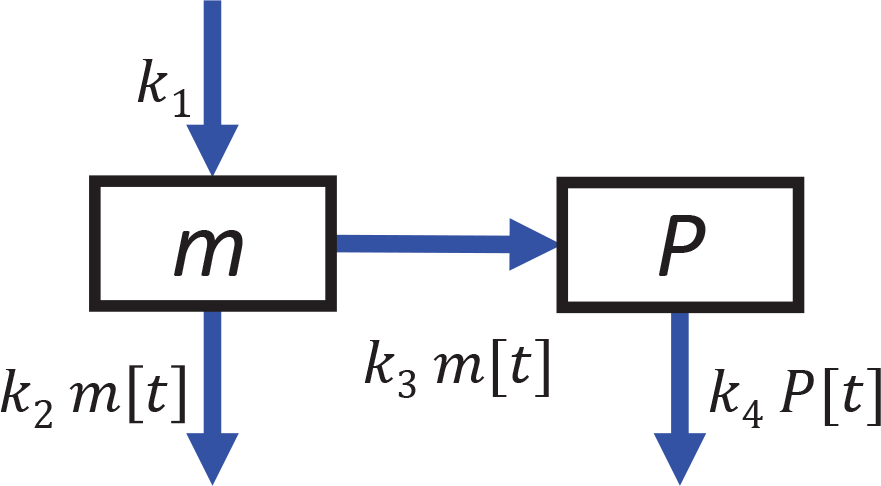
Linear model without feedback

The model has two species *m* and *P* and four rate constants *k*_1_*…k*_4_, as shown in Fig 4. *k*_1_ gives the rate of production of *m* via transcription of the gene for the protein *P*; *k*_2_ is a rate constant for degradation of *m*; *k*_3_ is the rate constant for translation of the mRNA into *P*; *k*_4_ is the rate constant for degradation of *P*. As we shall see, Hargrove and Schmidt did not plot the two-dimensional dynamics in a phase space “portrait”. The relative degradation rates of *m* and *P* turn out, perhaps surprisingly, to be a key determinant of the type of dynamics that is obtained. The kinetic equations are

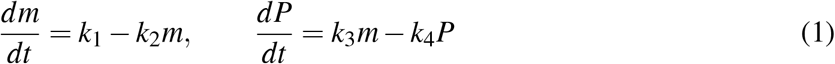

There is a single steady state (SS) solution when the net change rates of both species are zero:

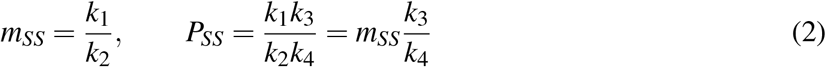

This single steady state is the basic target of the control system – we will verify later that the addition of feedback, though nonlinear, does not result in bifurcations to more steady states. Our model begins by specifying *m*_*SS*_, *P*_*SS*_ i.e. the parameter ratios in Eq. 2.

Next, we compute the dynamics around the steady state on a diagram of *normalized* concentrations *m/m*_*SS*_, *P/P*_*SS*_, with the steady state concentrations at (1, 1). We choose values of the rate constants at will to give a particular instantiation of the model. As the first example, we pick *k*_1_ = 20*/*17*, k*_2_ = 10*, k*_3_ = 34*, k*_4_ = 1. These parameters can be found as the first set in Table 1. The flow in the phase space of (*m/m*_*SS*_, *P/P*_*SS*_) is shown in Fig. 5(a). We see something very interesting. There is a “vertical structure” in the flow toward the steady state. There is first a general nearly horizontal fast flow toward the vertical. Then, flow takes place much more slowly asymptotically to the vertical toward the steady state. This latter flow is said to be along the vertical “slow manifold”.^15,16^ This can be understood in terms of the “linearized flow” near the SS in standard nonlinear dynamical analysis using the Jacobian; we go into detail in the Appendix. Dynamically, this behavior stems from the fact that the mRNA turnover rate is much faster than the protein turnover rate (*k*_2_*/k*_4_ ≫; 1). As a result, along the fast manifold, the mRNA concentration quickly reaches the steady state value *k*_1_*/k*_2_ and remains so, followed by the relatively slower change in *P*. This kind of dynamics can be described by the “quasi-steady state approximation” in traditional chemical kinetics.^13^

Investigating various parameter sets, we find that there are three general patterns of flow, which we designate in Fig. 5 as “vertical,” “focus,” and “diagonal.” The corresponding parameters are listed in the upper left part designated “linear” of Table 1. All of the flows have a separatrix along the vertical at *m* = *m*_*SS*_, resulting from a vertical eigenvector associated with the Jacobian matrix, as discussed in the Appendix. (In the vertical case, Fig. 5a, the separatrix corresponds to the slow manifold.) The patterns correspond to the following relations among the kinetic parameters:

1. Vertical case in Fig. 5a, *k*_2_ ≫ *k*_4_, (*k*_2_*/k*_4_ = 10 from Table 1): Fast relaxation of *m* (horizontal trajectories) to the steady state value, followed by slow change in protein concentration (vertical slow manifold). As mentioned earlier, this corresponds to the “quasi-steady state approximation” of traditional kinetics. Rosenfeld et al.^14^ made use of this approximation for the translational regulation case.
2. Focus case in Fig. 5b, *k*_2_ ≈ *k*_4_ (*k*_2_*/k*_4_ = 1 from Table 1) The eigenvalues −*k*_2_, −*k*_4_ are comparable, the trajectories are spiral-like so there is no separation in timescale into fast and slow manifolds. A similar case had been considered by Novák and Tyson.^17^
3. Diagonal case in Fig. 5c, *k*_2_ ≪ *k*_4_, (*k*_2_*/k*_4_ = 0.1 from Table 1): Here, the mRNA turnover rate *k*_2_ is much *slower* than the protein turnover rate *k*_4_.

It is very interesting that these basic patterns of the normalized phase portraits are determined by the ratio *k*_2_*/k*_4_ of *degradation rates* of the mRNA and the protein. The importance of the degradation rates was emphasized by Hargrove and Schmidt.^11^ (We will see in what follows that these three basic patterns play a crucial role in the classification of dynamics that the control system can exhibit.) Further, it turns out that the normalized (*m*_*SS*_, *P*_*SS*_) phase flows are a universal property of the linear model that depend only on the ratio *k*_2_*/k*_4_, invariant under any change in the other three variables in the parameter space. To understand this property mathematically, note that the linear partial differential equations in (1) can be solved analytically:

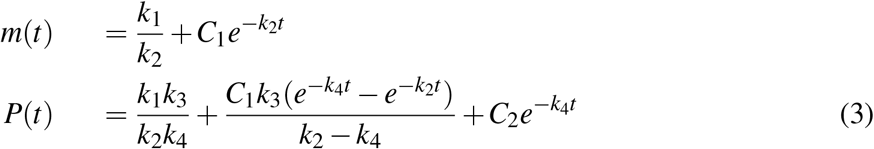

These depend only on *k*_2_, *k*_4_. This has the very important consequence that *the normalized phase flow depends only on the ratio k*_2_*/k*_4_. When, as we will consider later, the initial condition satisfies *m*[0] = *m_SS_*, we have *C*_1_ = 0*, m*(*t*) = *m_SS_*. Then the time-dependent protein concentration can be simplified as:

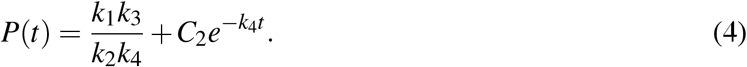

Clearly, in this special case, *P*(*t*) temporally depends only on *k*_4_.

**Figure 5:**
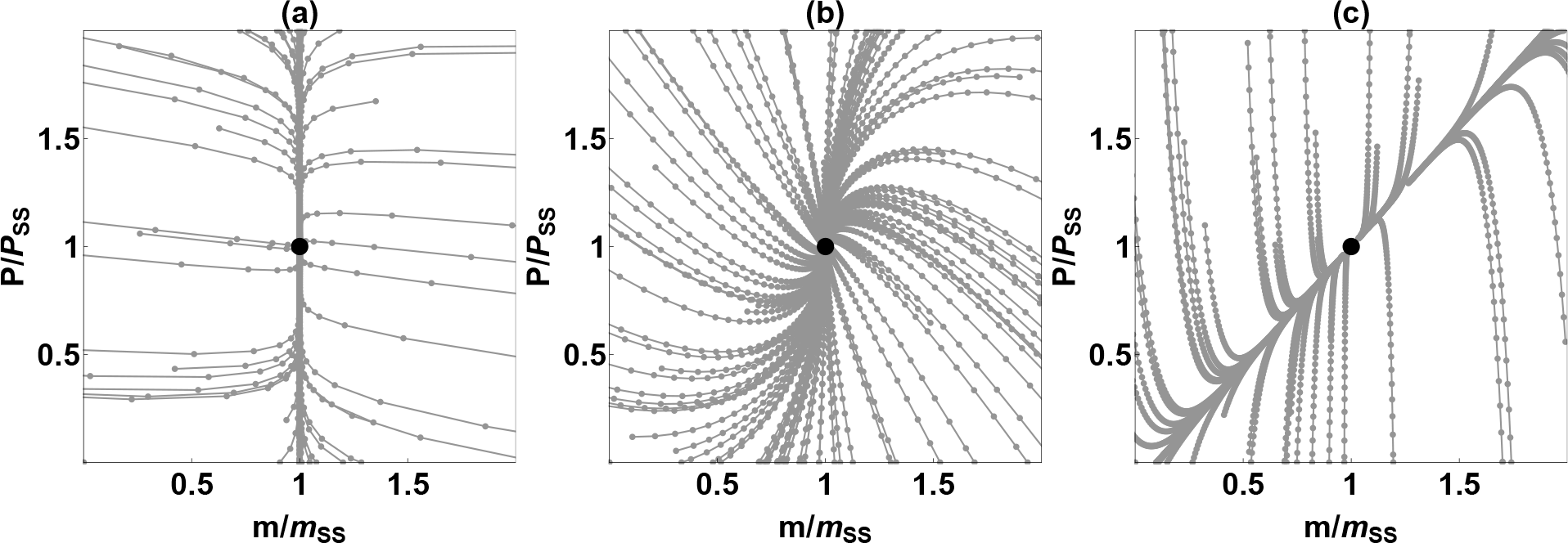
Representative normalized phase portraits for (a) vertical, (b) focus, and (c) diagonal types. The time step (space between the dots) is the same for all panels.

## Adding Feedback and cooperativity to tune the autoregulatory system

Already, these linear models constitute basic control schemes: in each, there is a steady state that is an attractor and the desired target for the dynamical system. However, this is a very primitive kind of control. Only the vertical profile has fast control directly to the *P*_*SS*_. The focus profile has wandering trajectories with arcing excursions. Even worse, the diagonal profile quickly directs the trajectory to the diagonal manifold, which however does not in general have *P* near the steady state value, so the trajectory gets stuck away from the target for a relatively long time. Moreover, the possibility, such as it is in each case, of achieving quick protein control by speeding up all the rates, also is extravagantly wasteful. After building *m* and *P* at great energetic cost, they are degraded, again at great cost! It is like filling buckets – one for *m*, a second, hydraulically linked one for *P* – that have holes in them designed to regulate the level in each bucket. The holes are the degradation processes with rate constants *k*_2_ and *k*_4_. To speed the system by a factor α, it suffices to increase all the parameters by α. This is certainly effective, but the metabolic cost just as certainly is extravagant. To switch metaphors, the processes of production of *m* and *P* are like an accelerator on a car, and the processes of degradation are like brakes. It is desirable to have a more subtle and discriminating system of accelerator and brakes. This is achieved by adding the element of nonlinear feedback with cooperativity. We will demonstrate these statements in the rest of the paper.

**Figure 6:**
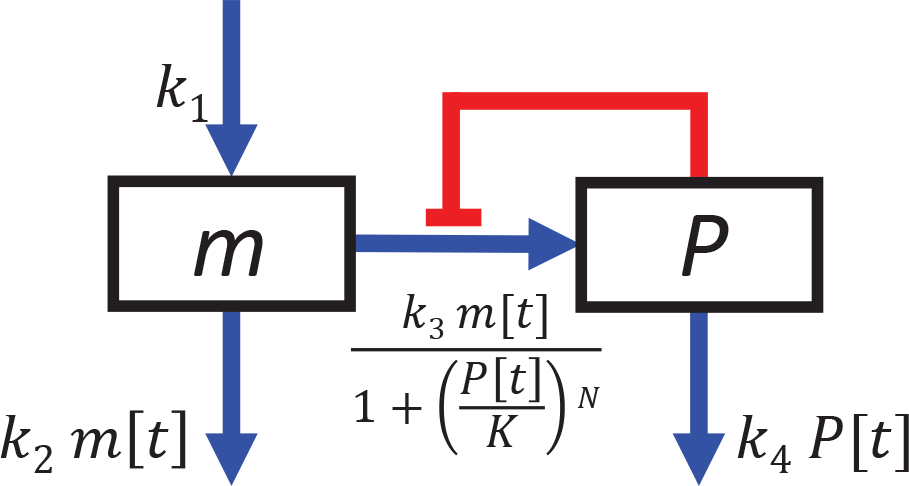
Schematic of autoregulation model with feedback and cooperativity, with rate constants and Hill-type factor.

We adopt a model with typical features of feedback and control, shown in Fig. 6, modifying the protein synthesis rate using “Hill-type” parameters often associated^9^ with feedback *K* and cooperativity *N*:

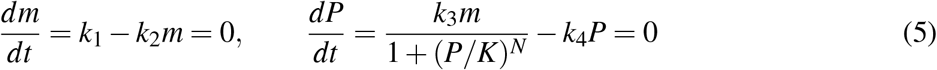

The denominator in the *P* rate equation is the feedback factor *X*, a function of *P* given by

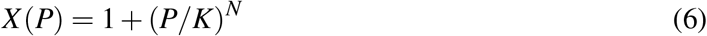

and is intended to represent the jamming of the mRNA by binding to its product protein *P*. The factor *X* (*P*) is a typical Hill-type expression involving the feedback parameter *K* and the cooperativity parameter *N*. We regard this kind of term in the way it is often used, as a phenomenological or empirical expression, not a literal expression in terms of a binding parameter *K* and cooperativity number *N*. The origin in enzyme kinetics and expanded empirical use of the Hill-type expressions is discussed in great detail by Ingalls.^9^

In effect, the concentration of available *m* is reduced by the factor 1*/X* (*P*). To represent this in the model, first we pick a value *X = X* (*P_SS_*) at the steady state. This modifies the expression for *P*_*SS*_ from Eq. 2 to

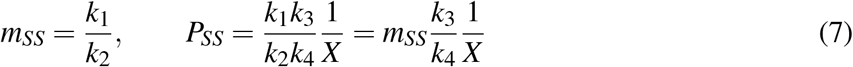

To maintain the same value of *P*_*SS*_ at the steady state, which we take to be the “goal” of the organism in “designing” the control scheme, with or without feedback, we need to compensate for 1*/X* (*P*) in equations (5,6). To do this, we choose to enhance the gene transcription rate constant *k*_1_ by the factor *X* – this seems the most likely of many possible scenarios involving the *k_i_*. This increases *m_SS_* from its value in the linear model by the factor *X*, to

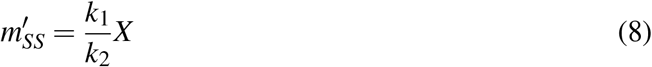

In effect, to maintain *P*_*SS*_ with the feedback, the system maintains a “reserve army” of messenger RNA via the enhanced rate constant *k*_1_.

## Systematic analysis of the autoregulatory system

So far we have established the basic framework of the linear system, with classification of the dynamics into three types based on the key ratio *k*_2_*/k*_4_; and then built in feedback with cooperativity. Now we systematically analyze the behavior and autoregulatory performance of the system, showing that with feedback there generally is far better control than with the corresponding linear system, with better time response to perturbation of the steady state while limiting the metabolic cost. We will do this in three steps: (A) Introduce prototypical individual trajectories from the phase space portraits, selected as being especially important examples of the autoregulatory process. (B) Then, consider systematic variation of the parameters in the kinetic model, in particular, of the key ratio *k*_2_*/k*_4_ with *k*_4_ fixed; and variation of this ratio with *k*_2_ fixed. (C) Examine the crucial indicator of autoregulatory efficacy, the time response of the protein concentration against various perturbations from the steady state concentrations *m_SS_, P_SS_* – first for the linear systems, and then the corresponding systems with feedback. We will see that the feedback mechanism gives greatly improved autoregulation. We present the tabular and visual content in Tables 1,2 and in Figs. 7,8, to which we refer repeatedly in the following.

### Prototype Trajectories

We will pay particular attention to trajectories of likely special importance. These are color coded in the figures. One type of trajectory is where the protein concentration is perturbed from its SS value, while the mRNA concentration is unchanged. An important example might be when the concentration of a particular protein is deliberately reduced in cancer chemotherapy.^18,19^ The purple and green trajectories in the figures have the protein concentration *P* displaced below and above the SS value, while *m* is kept at its steady state value, i.e. these trajectories start at (*m, P*) = (1, 2) and (1, 0) and end at the steady state (1, 1). Another important type of trajectory likely is where both *m* and *P* start at zero concentration, i.e. the system is “turning on.” This is seen in the orange trajectories, which start at (*m, P*) = (0,0). Finally, the cyan and blue trajectories correspond to perturbation in mRNA concentration, which could occur either naturally or due to the introduction of mRNA-binding species.^20^

### Systematics: Variation of Degradation Rates of mRNA and Protein

In Tables I, II and Figs. 7,8 we examine the effects of systematic variation in the kinetic parameters. We want to see how the dynamics change as the system changes – i.e. as the instantiation of our model defined by its particular parameters varies. We have supposed that in adding feedback to a linear model, e.g. through evolution, a primary criterion for an organism might be to preserve the SS protein concentration. Hence, we keep the steady state concentration *P*_*SS*_ = 4 in all cases. We consider systematic variation of the parameters *k*_2_, *k*_4_ for degradation of *m* and *P* since these seem to be the key to the pattern of the dynamics. In particular, we vary the key ratio *k*_2_*/k*_4_, first with *k*_4_ fixed; then with *k*_2_ fixed. We consider linear models, without feedback; and then corresponding models with feedback added. We will find that these parameter variations suffice to tell us most of what we want to know about the systematic behavior of the control systems.

In the linear models in Fig. 7a-c, we vary *k*_2_*/k*_4_ by keeping *k*_4_ fixed while varying *k*_2_ along with *k*_1_, and keeping *k*_3_ fixed. This preserves the value of the steady state concentrations *m_SS_, P_SS_*, according to Eqs. (1–2). See the corresponding values in the top part of Table I. In the linear models in Fig. 8a-c, we vary *k*_2_*/k*_4_ by keeping *k*_2_ fixed along with *k*_1_, while varying *k*_4_ along with *k*_3_. This again preserves *m*_*SS*_, *P*_*SS*_. The corresponding values are given in Table II.

In the feedback models in Fig. 7d-f, we use a feedback strength of *X*_*SS*_ = 17. To compensate for this, according to Section, we increase *k*_1_ from the corresponding linear models by a factor of 17. As in the case of linear models in Fig. 7a-c, we again vary *k*_2_*/k*_4_ by keeping *k*_4_ fixed, while varying *k*_2_ along with *k*_1_. This again preserves the value of the steady state concentrations *P*_*SS*_ (while changing the value of *m*_*SS*_ in the feedback models). The same parameter adjustment was implemented for Fig. 8d-f, except that now *k*_2_ and *k*_1_ are fixed, while *k*_4_ along with *k*_3_ are varied.

These systematic variations become clearer with perusal of the tables.

**Table 1:** Parameters used for Fig. 7. In all cases, *k*_3_ and *k*_4_ **are fixed; and** *k*_1_ **varies between linear and feedback systems by the factor** *X*_*SS*_.

|  | vertical, linear | focus, linear | diagonal, linear |
| --- | --- | --- | --- |
| $k_1$ | $\frac{20}{17}$ | $\frac{2}{17}$ | $\frac{2}{170}$ |
| $k_2$ | 10 | 1 | $\frac{1}{10}$ |
| $k_3$ | 34 | 34 | 34 |
| $k_4$ | 1 | 1 | 1 |
| $K$ | - | - | - |
| $N$ | - | - | - |
| $X_{SS}$ | 1 | 1 | 1 |
| $m_{SS}$ | $\frac{2}{17}$ | $\frac{2}{17}$ | $\frac{2}{17}$ |
| $P_{SS}$ | 4 | 4 | 4 |
| $k_2/k_4$ | 10 | 1 | $\frac{1}{10}$ |
| $\tau_1, \tau_2, \tau_3$ | 0.693, 0.693, 0.798 | 0.693, 0.693, 1.68 | 0.693, 0.693, 7.98 |
|  | vertical, feedback | focus, feedback | diagonal, feedback |
| $k_1$ | 20 | 2 | $\frac{2}{10}$ |
| $k_2$ | 10 | 1 | $\frac{1}{10}$ |
| $k_3$ | 34 | 34 | 34 |
| $k_4$ | 1 | 1 | 1 |
| $K$ | 4 | 4 | 4 |
| $N$ | 2 | 2 | 2 |
| $X_{SS}$ | 17 | 17 | 17 |
| $m_{SS}$ | 2 | 2 | 2 |
| $P_{SS}$ | 4 | 4 | 4 |
| $k_2/k_4$ | 10 | 1 | $\frac{1}{10}$ |
| $\tau_1, \tau_2, \tau_3$ | 0.311, 0.0361, 0.100 | 0.311, 0.0361, 0.297 | 0.311, 0.0361, 1.06 |
| $\tau_{linear}/\tau_{feedback}$ | 2.23, 19.18, 7.97 | 2.23, 19.18, 5.65 | 2.23, 19.18, 7.56 |

**Figure 7:**
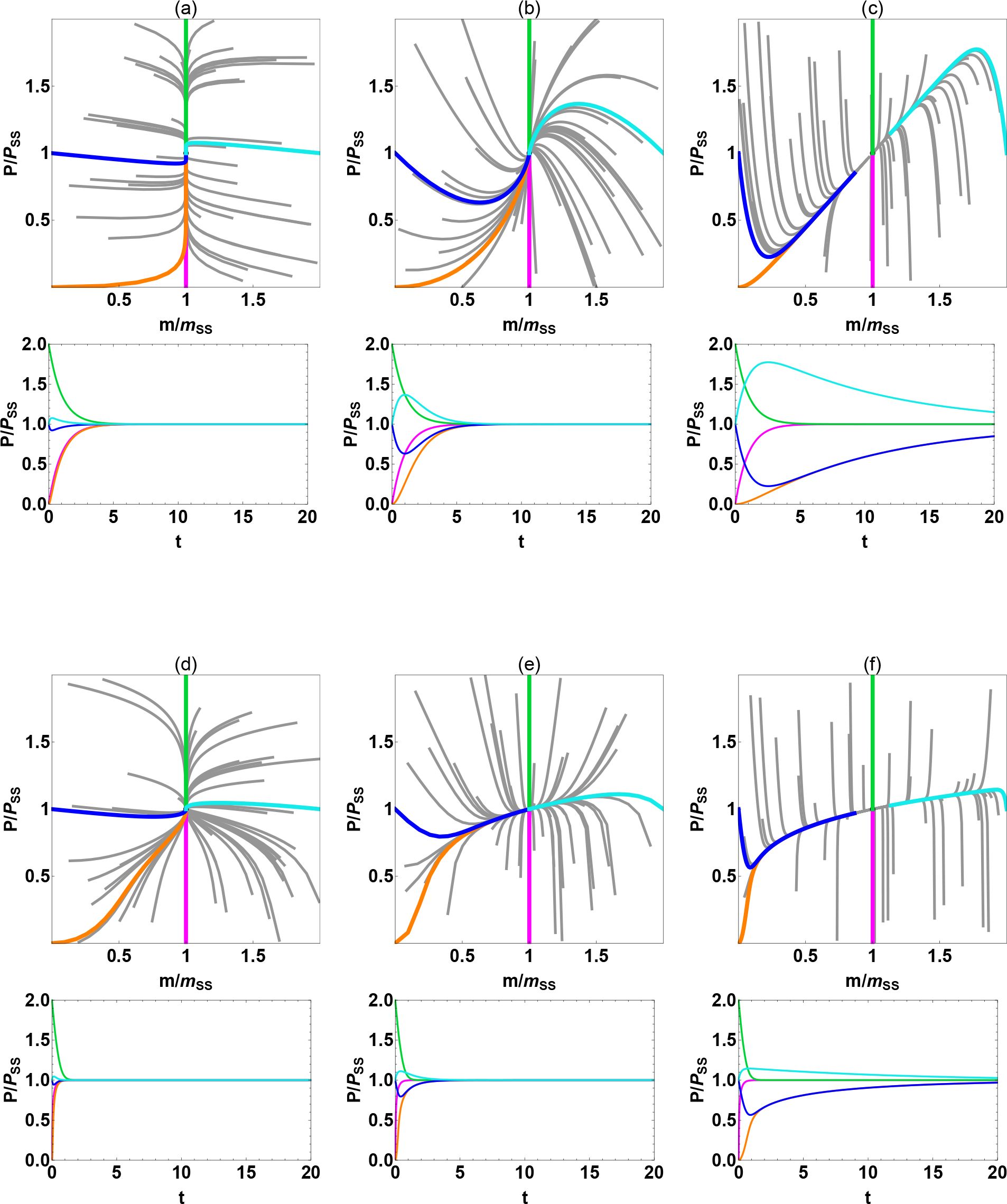
Normalized phase portraits and timecourses for *P/P_SS_* for Table I.

**Table 2:** Parameters used for Fig. 8. In all cases, *k*_2_ is fixed; *k*_1_ varies between linear and feedback systems by the factor *X_SS_*.

|  | vertical, linear | focus, linear | diagonal, linear |
| --- | --- | --- | --- |
| $k_1$ | $\frac{2}{17}$ | $\frac{2}{17}$ | $\frac{2}{17}$ |
| $k_2$ | 1 | 1 | 1 |
| $k_3$ | 3.4 | 34 | 340 |
| $k_4$ | 0.1 | 1 | 10 |
| $K$ | - | - | - |
| $N$ | - | - | - |
| $X_{SS}$ | 1 | 1 | 1 |
| $m_{SS}$ | $\frac{2}{17}$ | $\frac{2}{17}$ | $\frac{2}{17}$ |
| $P_{SS}$ | 4 | 4 | 4 |
| $k_2/k_4$ | 10 | 1 | $\frac{1}{10}$ |
| $\tau_1, \tau_2, \tau_3$ | 6.93, 6.93, 7.98 | 0.693, 0.693, 1.68 | 0.0693, 0.0693, 0.798 |
|  | vertical, feedback | focus, feedback | diagonal, feedback |
| $k_1$ | 2 | 2 | 2 |
| $k_2$ | 1 | 1 | 1 |
| $k_3$ | 3.4 | 34 | 340 |
| $k_4$ | 0.1 | 1 | 10 |
| $K$ | 4 | 4 | 4 |
| $N$ | 2 | 2 | 2 |
| $X_{SS}$ | 17 | 17 | 17 |
| $m_{SS}$ | 2 | 2 | 2 |
| $P_{SS}$ | 4 | 4 | 4 |
| $k_2/k_4$ | 10 | 1 | $\frac{1}{10}$ |
| $\tau_1, \tau_2, \tau_3$ | 3.11, 0.361, 1.00 | 0.311, 0.0361, 0.297 | 0.0311, 0.00361, 0.106 |
| $\tau_{linear}/\tau_{feedback}$ | 2.23, 19.18, 7.97 | 2.23, 19.18, 5.65 | 2.23, 19.18, 7.56 |

**Figure 8:**
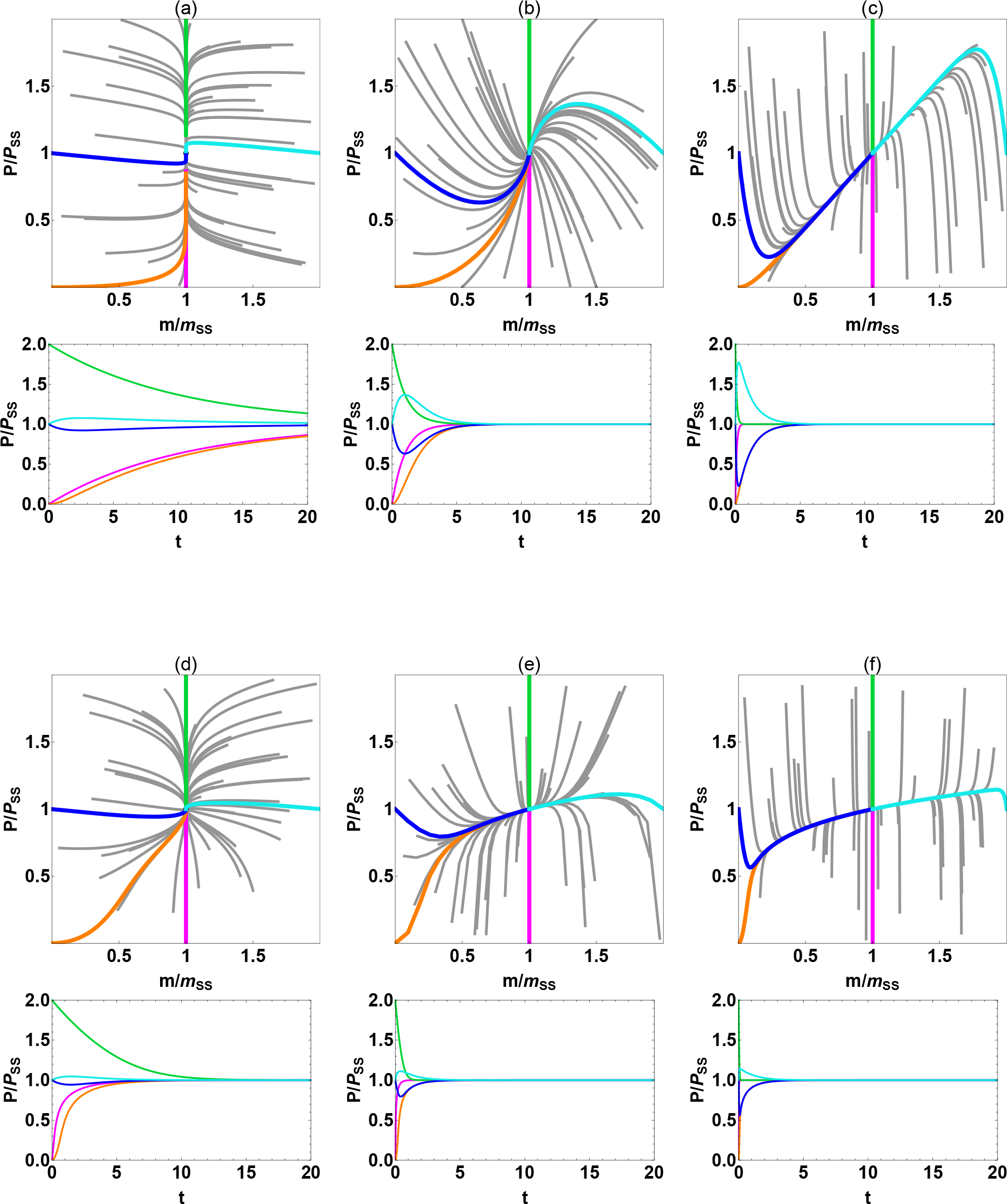
Normalized phase portraits and timecources for *P/P*_*SS*_ for Table II.

The computational dynamics are shown in Figs. 7, 8, each figure showing the phase portraits for the linear systems with color-coded trajectories (top row), the time response of the *P* concentration for various initial perturbations (second row), the phase portraits for the systems with feedback (third row), and the time response for the systems with feedback (fourth row). It is immediately evident that the phase portraits in all cases depend crucially on the ratio of degradation rates *k*_2_*/k*_4_. It is also immediately clear that there are similarities – in fact, exact invariances – between corresponding phase portraits, linear and feedback, in Figs. 7, 8. By invariance we mean equivalence between the dynamical flow throughout the normalized portrait.

The invariance of linear phase portraits between Figs. 7, 8 is easily explained by the fact that the time-dependent dynamics in the linear systems depends only on *k*_2_ and *k*_4_, as seen in Eq. 3. The explanation for the invariance evident in the feedback phase portraits between Figs. 7, 8 is not so evident, since by from Abel’s Impossibility Theorem,^21^ we do not have exact analytic solutions, for either the concentrations *P, m* or their derivatives in the rate equations, from which to make an invariance argument. The surprise disappears when one examines the corresponding parameter sets in Tables I, II. The invariances are between systems in which all the parameters *k*_1_…*k*_4_ are multiplied by a common factor of 10 or 1/10, i.e. in which there is merely an overall speedup or slowdown that obviously leaves the normalized phase portrait invariant. This common scaling factor, obtained following the procedure of parameter variations outlined above, is due to the constraints we have applied on *P*_*SS*_ and *m*_*SS*_, together with the relations in Eqs. 2, 7.

However, there is a further, genuine invariance puzzle that we will just touch upon here. As noted already, the linear system is invariant under any parameter change that maintains *k*_2_*/k*_4_. A surprising fact we have found is that the feedback system is also invariant under any parameter change that maintains both *k*_2_*/k*_4_ and *k*_1_*/k*_2_ × *k*_3_*/k*_4_. We will not discuss this much further here, as we do not yet completely understand it. We find that when we do vary *k*_1_*/k*_2_ *× k*_3_*/k*_4_, the perturbation to the phase portrait is rather mild. This “symmetry” of the kinetic equations clearly merits future exploration. In fact, these invariance and near-invariance properties are an important prediction of our model that could be tested in real biological systems with translational autoregulation.

### Autoregulatory power: Time response of the linear and feedback systems

Now we will examine the representative individual trajectories in Figs. 7, 8 with a view toward sizing up their powers of autoregulation. The key criteria for regulation have to do with the response of the system under a perturbation – of whatever cause – foremost, the behavior of the protein concentration *P*; secondarily, the mRNA concentration *m*; with consideration of time and metabolic factors. We will often make reference to the color-coding of the trajectories in the figures.

We focus first on the three phase space portraits of the linear system in Fig. 7(a)-(c), associated with the parameters in Table 1; then on the three portraits of the feedback system in (d)-(f). For each of the three systems in each row, we have chosen trajectories corresponding to perturbed conditions that might be of prime importance, as touched upon in Section. To reiterate from Section: the green trajectory in each figure starts with an excess of *P* with *m* fixed at the SS value. The purple trajectory starts at reduced value of *P* with *m* fixed at its SS value. The orange trajectory is for starting the system at zero values of both *m* and *P*. All of these trajectories eventually return to the SS. Also shown are trajectories (cyan, blue) in which the system is perturbed to deficient or excess values of *m* with *P* fixed at its SS value. The time response of the *P* concentration for the various trajectories is shown under the phase portraits in the figures, in rows 2 and 4.

The green and purple “vertical trajectories” are each identical among the three linear systems and also among the feedback systems (with different response rates than the linear systems). This is because the mRNA concentration is fixed at *m_SS_* along these trajectories, as follows mathematically from Eq. 4. Moreover, the green and purple trajectories in each portrait have the same time dependence, again from Eq. 4. On the other hand, the other trajectories that have *m* dependence differ greatly among the linear systems, and also among the feedback systems, showing the importance of the full two-species dynamics associated with the three classes and *k*_2_*/k*_4_. It is evident from the time dependence of *P* for the various trajectories, shown in the second and fourth rows, that with feedback there is a great speedup in response times. It is noteworthy that the purple and green trajectories differ in each feedback system portrait – the green (excess *P*) is much slower than the purple (deficit of *P*). We will examine the time response in more detail later shortly.

Fig. 8 shows the corresponding information for the systems in Table II where *k*_4_ is varied, instead of *k*_2_. Not all of the statements regarding Fig. 7 apply in these cases. Now the protein rate constants *k*_3_*, k*_4_ vary, and so do the time responses across rows 2 and 4 – despite the exact invariance of the phase portraits between the two figures. There is again a great speedup in the feedback systems. Examination of the figures is probably more illuminating than further verbal description.

We now turn to a quantitative measure of the response times and autoregulatory power of the various systems. We take this, where it applies, to be the time τ*_i_* for the *P* concentration along a given trajectory to return halfway from its starting point to *P_SS_*. We give the values of τ_1_…τ_3_ for the green, purple, and orange trajectories respectively at the bottom half of Tables I, II, together with the ratio of corresponding τ_*linear*_/τ_*feedback*_. This ratio is an indication of response speedup due to the feedback; a larger ratio indicates a greater speedup. A lot of regularities are evident among the τ*_i_*. These are related to the symmetries, invariances, and parameter ratios described above among the parameters and phase portraits, e.g. factors of 10.

We can see the great advantages to response time afforded by feedback as compared to the crude method in the linear system of just multiplying all the parameters by a common factor (thus providing an overall speedup – at great metabolic cost). Fig. 8a compared to Fig. 7a shows the result of multiplying all the rates by a common factor. Comparing Fig. 8a to Fig. 8d shows that a generally comparable speedup is obtained with feedback, by changing only one of the parameters *k*_1_, the rate of transcription of the DNA, instead of all of them. This should be a great metabolic advantage, since the number of *m* and especially of *P* molecules produced by just one gene can be very large, ranging in the thousands.^12,22^ Similar observations pertain to Fig. 7c compared to Fig. 8c; and then Fig. 7c compared to Fig. 7d. The secret of the feedback regulation efficiency is basically that feedback, at relatively little cost, replaces degradation as the control mechanism, while sacrificing little in speed. The “brakes” are applied selectively, only as needed.

There is an interesting partial exception to these statements. Note that the green trajectories (relieving excess *P*) in the feedback systems show a much slower response rate than the purple trajectories (relieving deficient *P*). This might seem surprising in that the control scheme depicted in Fig. 1 is based on jamming the production of *P*. From the computed dynamics, apparently, greater inhibition of *P* production at excess concentrations is more effective than lesser inhibition at deficient concentrations.

## Discussion and conclusions

We end with a summing up, and a prospectus for future application of kinetic modeling of autoregulation to important biological systems and problems, such as chemotherapy of many cancers, briefly discussed below. We have developed a two-component kinetic model for translational autoregulation, with mRNA concentration *m* and protein concentration *P* as dynamical variables. The basic linear system gives a “barebones” model with regulation of the system toward the target steady state. However, the linear model is extravagantly wasteful: to speed up control against perturbations away from the steady state, multiple rate constants must be increased, generally at great metabolic cost. To attain more efficient control, feedback with cooperativity can be added to the model. Adding feedback with cooperativity dramatically improves the autoregulatory response. This attains the objective of control that is targeted and fast, yet efficient. In general, in comparison to the linear systems, which depend entirely on degradation of *m* and *P* for control, the feedback achieves comparable speedup in response time at much lower metabolic cost.

For perturbation of *P* alone, with *m* maintained at its steady state value, the system is basically one-dimensional, with correspondingly simple kinetic relations. However, with perturbation of both *m* and *P*, the full two-dimensional description is essential, with a great variety of dynamical possibilities in the three basic phase space structures. Massive reorganization of the dynamics can occur when the system parameters are changed. If the parameters become changed by the organism in response to a perturbation – e.g. if either or both of the degradation rates *k*_2_, *k*_4_ are changed – a new instance of the autoregulatory system is created which changes the entire behavior of the system.

To a large extent, the ratio of degradation rates *k*_2_*/k*_4_ for *m* and *P* governs the structure of the dynamical flow in the phase space portrait. This means that linear systems are exactly invariant under all parameter changes that preserve *k*_2_*/k*_4_. Systems with feedback are exactly invariant under parameter changes that preserve both *k*_2_*/k*_4_ and *k*_1_*/k*_2_ *× k*_3_*/k*_4_ – an interesting empirical “symmetry” that is still under investigation. Computation shows that systems with feedback change mildly under changes in *k*_1_*/k*_2_ *× k*_3_*/k*_4_ that preserve *k*_2_*/k*_4_. Consistent with the above statements, the autoregulatory response for both the linear and the feedback systems depends greatly on the phase space structure associated with *k*_2_*/k*_4_. These invariance properties are an important prediction of our model that possibly could be tested in real biological systems that have translational autoregulation.

Both transcriptional and translational regulation are a widespread phenomenon of genetic control.^3^ In future work we plan to investigate interesting contrasts in the kinetics and dynamics of these types of autoregulation. One of the pioneering examples of translational autoregulation involves the protein gp32 in replication of the virus T4 during infection of *E. coli*.^5^ We hope to analyze a specific example of our model built with parameters derived as much as possible from experiment.

This paradigmatic translational control mechanism has been found to be important in cells.^23–25^ An example appears to be autoregulation of thymidylate synthase (TS), an enzyme which is very important in both normal and cancerous cells, the latter including many of the most common cancers. The TS autoregulation is believed^26,27^ to be crucial in the development in cancer cells of resistance against chemotherapy drugs, many of which (e.g. 5-fluorouracil) specifically bind to TS in order to hinder DNA synthesis in cancer cells or induce apoptosis. This is very much like our purple trajectories in Figs. 7, 8. Then, it is possible that some reorganization takes place that would correspond to change in one or more of the parameters of our model, associated with the unfortunate development of resistance to the drug. We believe that building quantitative kinetic schemes tailored to biological systems is essential for thorough understanding of the chemotherapeutic process, and perhaps even intervening in new ways.

We have tried to convey that it is pretty much hopeless to fully apprehend the dynamics of even so simple a model as ours here without the quantitative analysis of the full phase space dynamics. This statement surely must carry over to more complex genetic regulatory networks – underlining how daunting is a full understanding of their behavior. This underlines the difficulty of realizing the promise of Waddington’s heuristic notion of the “epigenetic landscape”.^28–30^

As a final remark, we make the observation that the autoregulatory model developed here does not function much like a “program.” There is no logical scheme here, other than the tendency of the complete system to revert to the SS. The “intelligence” in the system is simply in the dynamics. If biological systems function something like an operating system running programs, it happens at a larger or more complex scale of organization than the simple system here – which would likely be a component of some such larger system.

## Appendix Jacobian analysis

Generally, for the differential equation set

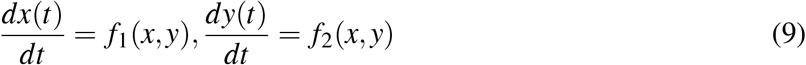

a steady state (*x*_0_*, y*_0_) is defined as a point in the phase space where *f*_1_ = *f*_2_ = 0. Near such a point, the dynamics can be approximated linearly as^31^:

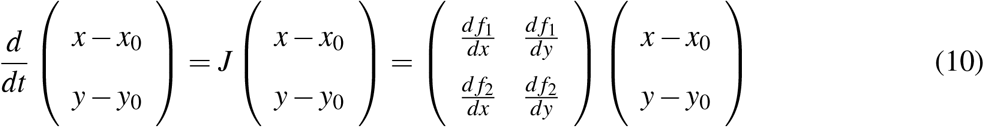

where (*x*(*t*) −*x*_0_*, y*(*t*) −*y*_0_) is a vector and *J* a matrix. Diagonalization of *J* yields two eigenvalues λ_1_, λ_2_ and their associated eigenvectors 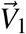,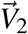. At a stable steady state, λ_1_ < 0 and λ_2_ < 0, and the solution of the linear system has the following form:^31^
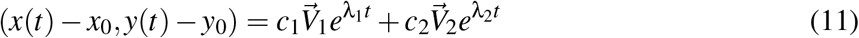

Hence, when |λ_1_*| ≫ |*λ_2_*|*, there is a natural separation of time scale, with the fast and slow components along the directions of 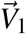 and 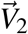, respectively. Note that the eigenvectors may not be orthogonal to each other when *J* is not symmetric, which is the case here. In the no-feedback model, from the kinetic equations (1) we obtain

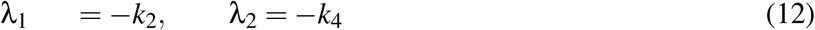

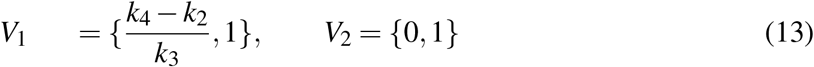

Note that in this case *V*_2_ always points vertically, since *dm/dt* does not depend on *P* (while *dP/dt* depends on *m*). This has the consequence that trajectories on left and right of the vertical are strictly separated, as is apparent from the phase space portraits. By considering different values of *k*_1_ − *k*_4_, the dynamics can be divided into the 3 categories depending on the ratio of *k*_2_*/k*_4_.

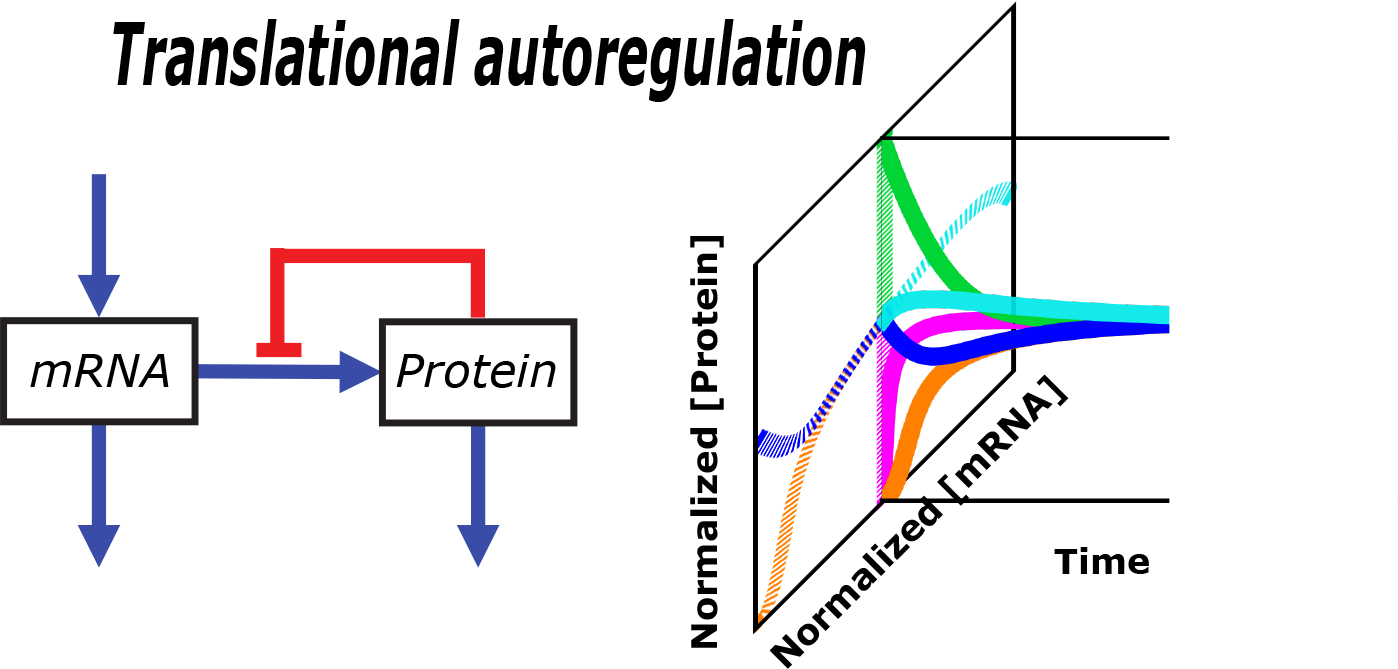

## Acknowledgement

We would like to thank Pete von Hippel and members of his group seminar for many stimulating discussions about translational autoregulation, especially of gp32 in the T4 - E. coli system.

